# Mu-opioid receptor activation potentiates excitatory transmission at the habenulo-peduncular synapse

**DOI:** 10.1101/2024.12.10.627842

**Authors:** Sarthak M. Singhal, Agata Szlaga, Yen-Chu Chen, William S. Conrad, Thomas S. Hnasko

## Abstract

The continuing opioid epidemic poses a huge burden on public health. Identifying the neurocircuitry involved and how opioids modulate their signaling is essential for developing new therapeutic strategies. The medial habenula (MHb) is a small epithalamic structure that projects predominantly to the interpeduncular nucleus (IPN) and represents a mu-opioid receptor (MOR) hotspot. This habenulo-peduncular (HP) circuit can regulate nicotine and opioid withdrawal; however, little is known about the physiological impact of MOR on its function. Using MOR-reporter mice, we observed that MORs are expressed in a subset of MHb and IPN cells. Patch-clamp recordings revealed that MOR activation inhibited action potential firing in MOR^+^ MHb neurons and induced an inhibitory outward current in IPN neurons, consistent with canonical inhibitory effects of MOR. We next used optogenetics to stimulate MOR^+^ MHb axons to investigate the effects of MOR activation on excitatory transmission at the HP synapse. In contrast to its inhibitory effects elsewhere, MOR activation significantly potentiated evoked glutamatergic transmission to IPN. The facilitatory effects of MOR activation on glutamate co-release was also observed from cholinergic-defined HP synapses. The potentiation of excitatory transmission mediated by MOR activation persisted in the presence of blockers of GABA receptors or voltage-gated sodium channels, suggesting a monosynaptic mechanism. Finally, disruption of MOR in the MHb abolished the faciliatory action of DAMGO, indicating that this non-canonical effect of MOR activation on excitatory neurotransmission at the HP synapse is dependent on pre-synaptic MOR expression. Our study demonstrates canonical inhibitory effects of MOR activation in somatodendritic compartments, but non-canonical faciliatory effects on evoked glutamate transmission at the HP synapse, establishing a new mode by which MOR can modulate neuronal function.

## Introduction

The opioid addiction crisis poses a huge burden on public health. Relapse commonly occurs after cessation of drug use, in part due to the onset of opioid withdrawal symptoms (Kosten and Baxter, 2019; Volkow et al., 2019). The locus coeruleus and mesolimbic system have been implicated in regulating opioid withdrawal symptoms (Pergolizzi et al., 2020; Welsch et al., 2020), however, mu-opioid receptors (MOR) are expressed widely. To open new avenues for therapeutic intervention there is a clear need to identify how other neural systems contribute.

The medial habenula (MHb) is a bilateral, epithalamic structure that is part of the habenular complex and expresses one of the highest densities of mu-opioid receptor (MOR) (Mansour et al., 1987; Kitchen et al., 1997; Gackenheimer et al., 2005; Gardon et al., 2014). It is primarily glutamatergic but also contains neurons that co-synthesize either acetylcholine (ACh) or the tachykinins, substance P and neurokinin B (Ren et al., 2011; Viswanath et al., 2013). The MHb receives inputs from different septal nuclei and sends dense projections to the interpeduncular nucleus (IPN) through the fasciculus retroflexus, comprising the habenulo-peduncular (HP) circuit (McLaughlin et al., 2017). MHb projections to IPN are topographically organized with dorsal MHb neurons projecting mainly to the rostral (IPR) and lateral (IPL) subregions of the IPN, and ventral MHb neurons sending projections to the IPR and central subregions (IPC) (Molas et al., 2017). The IPN is a midline midbrain nucleus composed primarily of GABAergic neurons and sends efferents to raphe and tegmental nuclei, amongst others (Lima et al., 2017; Quina et al., 2017).

The HP circuit is important in regulating nicotine dependence and withdrawal, and it is more generally implicated in the regulation of anxiety and mood (Salas et al., 2009; Antolin-Fontes et al., 2015; McLaughlin et al., 2017; Molas et al., 2017; Ables et al., 2023). For example, hyperactivity of the HP circuit increased depression-like behaviors in rats (Xu et al., 2018), and GABA_B_ and CB1 receptor actions at the HP synapse regulated fear behavior and the expression of aversive memory (Soria-Gómez et al., 2015; Zhang et al., 2016). Recent behavioral pharmacology studies also revealed a role for MOR expression in the MHb in opioid dependence. Deletion of MOR from β4 nicotinic receptor expressing cells, which are enriched in MHb, attenuated the affective and physical signs of morphine withdrawal (Boulos et al., 2020) and impaired social behaviors (Allain et al., 2022). Optogenetic stimulation of habenular MOR projections triggered avoidance and despair-like behavior (Bailly et al., 2023).

MORs are G_i_-linked G-protein coupled receptors (GPCRs), and their activation can inhibit cAMP production, open inward rectifying potassium channels (GIRKs), and thereby make neurons less excitable. MORs also localize to axon terminals where they can inhibit synaptic transmission (Williams et al., 2001; Al-Hasani and Bruchas, 2011; Che and Roth, 2023). However, it is unclear how MOR activation affects excitability or neurotransmission within the HP circuit. Understanding the consequences of MOR signaling in the HP circuit is a key step toward revealing its role in opioid addiction.

Here, we used reporter mice and *in situ* hybridization to visualize MOR expression in a subset of MHb and IPN neurons, and used patch-clamp electrophysiology to reveal how MOR activation influences MHb and IPN neurons and synapses. We found that MOR has canonical effects, inhibiting MOR-expressing neurons in both MHb and IPN. In contrast, we found that MOR activation markedly potentiated evoked excitatory transmission at the HP synapse, including from glutamate-releasing HP synapses made by cholinergic neurons. Finally, we provide pharmacological and genetic evidence indicating that that these facilitatory effects on neurotransmission are due to the direct activation of pre-synaptic MORs at the HP synapse. To our knowledge this is the first report of a direct faciliatory effect of MOR activation on neurotransmission, suggesting a new non-canonical mode of MOR signaling. In sum, we demonstrate compartment-specific effects of MOR activation on neuronal excitability and excitatory transmission in the HP circuit and provide the first electrophysiological characterization of the physiological consequences of MOR in this circuit which is of known importance to aversion, anxiety, and addiction.

## Results

### Mu-opioid receptor (MOR) is expressed in MHb and IPN neurons and its activation is inhibitory

To investigate the expression of MOR in neurons comprising the HP circuit we generated mice that express a fluorescent reporter in cells that expressed MOR (MOR-Cre). Within the MHb, reporter expression was observed in a subset of MHb neurons concentrated in lateral and dorsal subregions across its rostral to caudal extent (**Figure 1A**). We next used fluorescent *in situ* hybridization by RNAscope to validate this expression pattern. We used probes targeting *Oprm1* (MOR), as well as a glutamatergic marker (vesicular glutamate transporter 2, VGLUT2, *Slc17a6*), and a cholinergic marker (choline acetyltransferase, *Chat*). As expected, mRNA encoding MOR was concentrated in lateral and dorsal MHb subregions (**Supplemental Figure 1A**), where it partially colocalized with cholinergic and glutamatergic markers (**Supplemental Figure 1B**). We next used patch-clamp electrophysiology (cell-attached recordings) to test how MOR activation changed the firing rate of spontaneously active MHb neurons. When blind patching from un-tagged MHb neurons in wild-type mice, we found only a subset (4/10 neurons) appeared to slow firing when the MOR agonist DAMGO (5 µM) was applied (**Supplemental Figure 1C-D**) (p= 0.03, Friedman test). However, when recording from defined MOR^+^ neurons using MOR-Cre reporter mice (**Figure 1B**), DAMGO consistently and significantly inhibited MHb neuron firing, and this inhibition was reversed upon application of the MOR antagonist naloxone (**Figure 1C-D**) (p= 0.0003, Friedman test). These data show that MOR is expressed in a subset of MHb neurons where it has inhibitory effects on cell firing consistent with canonical actions of MOR activation.

**Figure 1.**
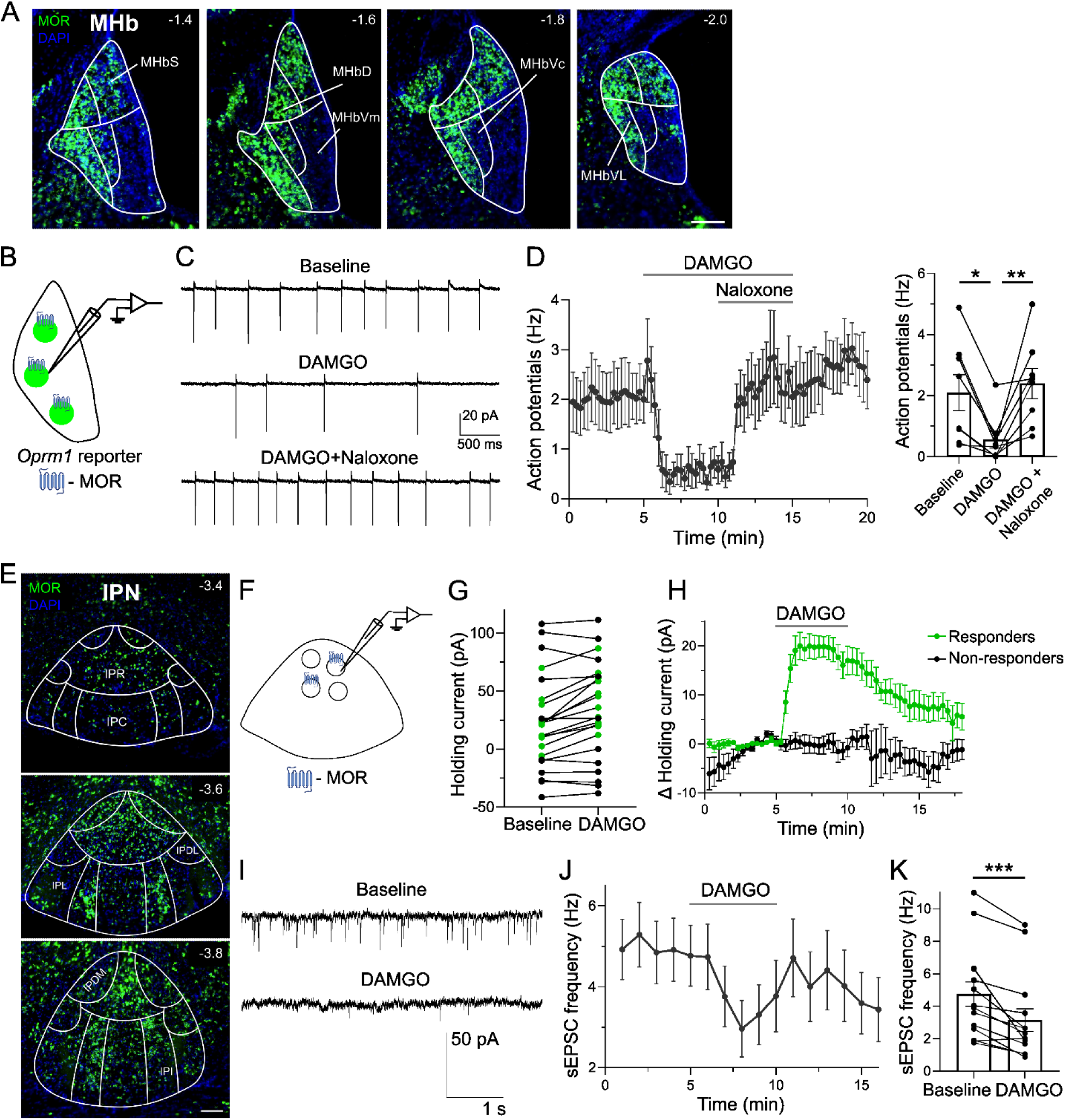
Inhibitory effects of MOR activation in the MHb and IPN. **A**) Example images through MHb show Zs-Green reporter expression (green) resulting from MOR-Cre; DAPI in blue. MHb subregions: S- superior, D- dorsal, Vm- ventral medial, Vc- ventral central, VL- ventral lateral; scale, 100 μm. **B**) Schematic of cell-attached recordings from MOR^+^ cells in the MHb obtained from ZsGreen reporter mice. **C**) Example recording and **D**) averaged time-trace (left) and bar-graph (right) of cell-attached recordings showing DAMGO (5 μM)-mediated inhibition of action-potentials which was reversed by naloxone (5 μM) application (n=8 neurons); Friedman test, *p< 0.05, **p< 0.01. **E**) Representative images through IPN show Zs-Green reporter expression (green) resulting from MOR-Cre; DAPI in blue. IPN subregions: IPR- rostral, IPC- caudal, IPL- lateral, IPDL- dorsal lateral, IPDM- dorsal medial, IPI- intermediate; scale, 100 μm. **F**) Schematic of whole-cell recordings from untagged IPN cells, some of which expressed MOR. **G**) DAMGO (1 μM) application increased holding current in a subset of IPN neurons (n=12/23), pseudo- colored green, consistent with post-synaptic expression of MOR in a subset of IPR neurons. **H**) Averaged time-traces show DAMGO-induced outward current in responding (n=12) versus non-responding neurons (n=11). **I**) Example recording, **J**) averaged time- trace, and **K**) bar-graph of whole-cell recordings showing DAMGO (1 μM)-mediated inhibition of sEPSC frequency in IPN neurons (n=14); Wilcoxon test, ***p< 0.001.

We next turned our attention to the IPN. Using both MOR-Cre reporter mice (**Figure 1E**) and RNAscope (**Supplemental Figure 1E)**, we found that a subset of IPN neurons expressed MOR. These cells were present across the rostral to caudal extent of IPN and showed notable density in rostral IPN (IPR) and intermediate IPN (IPI). To test how MOR activation influences IPN cells, whole-cell voltage-clamp recordings were made from untagged neurons in IPR (**Figure 1F**). We found that DAMGO application led to a steep increase in the holding current in a subset of recorded neurons (n= 12/23, **Figure 1G-H**). This suggests that in a subset of IPN neurons, presumably those expressing MOR, MOR activation recruits a hyperpolarizing, or inhibitory, conductance. We also measured the rate of spontaneous excitatory postsynaptic currents (sEPSCs) in IPR neurons. The frequency of sEPSCs was also significantly reduced by DAMGO application (**Figure 1I-K**) (p= 0.0002, Wilcoxon test), suggesting that spontaneous release from MOR-expressing excitatory inputs on to IPR neurons can be inhibited by MOR activation.

Overall, these data show that MOR is expressed in a subset of MHb and IPN neurons, and that at multiple locations within the HP circuit MOR activation can drive canonical inhibitory effects.

### MOR activation potentiates evoked glutamate neurotransmission at the HP synapse

Next, we investigated how MOR activation impacts evoked neurotransmission at the HP synapse. We injected AAV into the MHb of MOR-Cre mice (**Figure 2A**) to express ChR2:mCherry in MOR-expressing cells (**Figure 2B**) and terminals in the IPN, which were mainly observed in the IPR and IPL subregions (**Figure 2C**). Spread of ChR2:mCherry expression was typically seen in areas adjacent to MHb, including the lateral habenula (LHb) and paraventricular thalamus (PVT) (**Figure 2B**), however these adjacent regions project little or none to IPN (Quina et al., 2015; Lima et al., 2017). Following AAV injection, we made whole-cell patch-clamp (voltage-clamp) recordings from IPR neurons that were surrounded by ChR2:mCherry^+^ fibers. We delivered two pulses of blue light to assess fast excitatory post-synaptic currents that were evoked in response to optogenetic stimulation (oEPSCs). We found that bath application of DAMGO consistently and significantly increased oEPSC amplitude (**Figure 2D-F**) (t_6_= 4.1, p= 0.006). In a subset of cells (n=3), AMPA- and NMDA-type glutamate receptor blockers (DNQX and AP5) were applied after DAMGO washout, and these abolished the oEPSCs (**Figure 2G-H**), indicating they were dependent on glutamate release.

**Figure 2.**
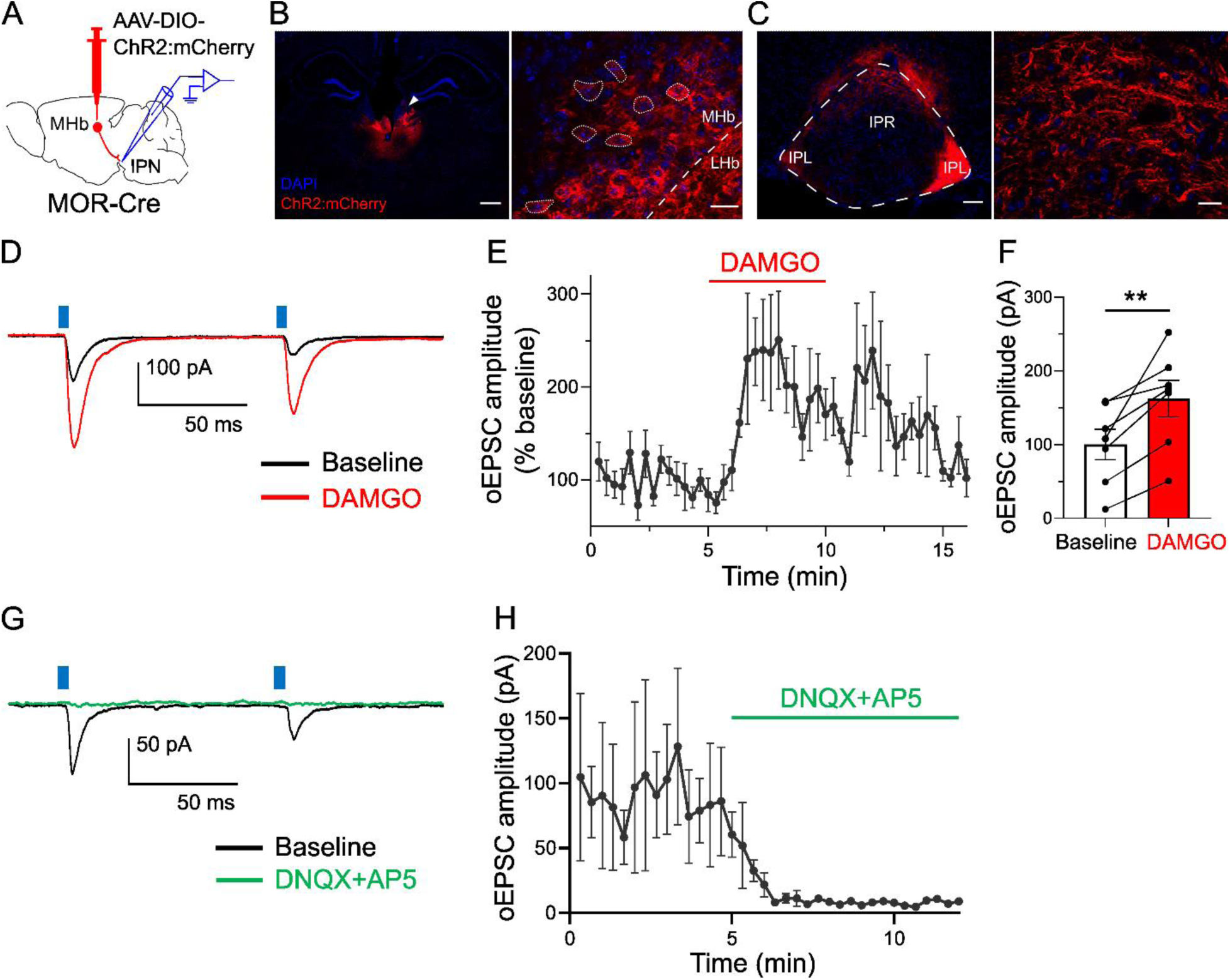
MOR activation enhances excitatory transmission at the HP synapse. **A**) Experiment design showing patch-clamp electrophysiology in the IPN following AAV injection into MHb of MOR-Cre mice. **B**) (left) Image of MHb injection site (white arrowhead); scale, 500 μm. (right) Dotted lines outline ChR2:mCherry^+^ cells in MHb; scale, 20 μm. **C**) (left) ChR2:mCherry^+^ projections in rostral (IPR) and lateral (IPL) IPN subregions, scale: 100 μm. (right) Magnified image of axonal fibers in IPR; scale, 20 μm. **D**) Traces from an example IPR neuron showing oEPSCs, before and after application of DAMGO (1 μM). **E**) Averaged time-trace and **F**) bar graph show DAMGO-mediated increase in oEPSC amplitude at the HP synapse (n=7); paired t-test, **p< 0.01. **G**) Example trace from a cell and **H**) averaged time trace show that application of glutamate receptor blockers, DNQX (10 μM) plus AP5 (40 μM) abolished oEPSCs (n=3).

We also tested the effects of MOR activation on evoked neurotransmission in IPN sections obtained from crossing MOR-Cre to Ai27 or Ai32 mice, which results in Cre-dependent expression of ChR2:mCherry or ChR2:YFP in cells that expressed MOR. Similar to the AAV-based approach, we found that DAMGO led to an increase in oEPSC size in IPR neurons (**Supplemental Figure 2A-D**) (t_4_= 3.1, p= 0.035). However, in these experiments we observed low rates of connectivity, with only 8 of 41 patched IPR neurons showing oEPSCs. Moreover, in 17/41 IPR neurons we detected fast photocurrents with an onset latency of <1ms, visible in the example trace (**Supplemental Figure 2B**). These photocurrents are consistent with a subset of IPR neurons expressing MOR, but due to these complicating factors we discontinued use of this preparation.

Overall, our results indicate that MOR activation can result in a non-canonical potentiation of evoked glutamatergic transmission at the HP synapse.

### MOR potentiates excitatory neurotransmission at the HP synapse through a monosynaptic mechanism

The non-canonical potentiating effect of MOR activation on excitatory neurotransmission that we observed at the HP circuit was surprising; to our knowledge, such an effect has never been reported before for MOR. We therefore began a series of studies to investigate the mechanism further. We first aimed to understand whether the oEPSC potentiation might be due to recruitment of a polysynaptic circuit. IPR neurons are primarily GABAergic and subsets of IPR neurons express MOR, thus MOR activation on IPN neurons could lead to changes in GABA signaling that contribute to oEPSC potentiation. To test if the effects of MOR activation on excitatory neurotransmission at the HP synapse are dependent on indirect effects through local GABA signaling, we tested the effects of DAMGO in the presence of GABA_A_ and GABA_B_ receptor blockers gabazine and CGP-55845, at pharmacologically validated concentrations (**Supplemental Figure 3A and 4E**). Following pre-treatment with either gabazine+CGP or vehicle, bath application of DAMGO increased the oEPSC amplitude at the HP synapse (**Figure 3A-B**) (main effect of treatment: F_1,12_= 13.4, p= 0.003; group x treatment interaction: F_1,12_= 3.3, p= 0.09). Notably, we observed a DAMGO-mediated inhibition in oEPSC amplitude in 2 neurons in this experiment (see further below), both happened to be in the vehicle group (**Figure 3B**). Excitatory effects of DAMGO on neurotransmission were similarly observed in a separate group of neurons pre-treated with the GABA_A_ receptor blocker picrotoxin (PTX) (**Supplemental Figure 3B-C**) (main effect of treatment: F_1,5_= 13.9, p= 0.01; group x treatment interaction: F_1,5_= 0.7, p= 0.44). These results indicate that DAMGO-mediated potentiation of excitatory neurotransmission at the HP synapse persists in the presence of GABA receptor blockade and are therefore not dependent on recruitment of local GABA signaling mechanisms.

**Figure 3.**
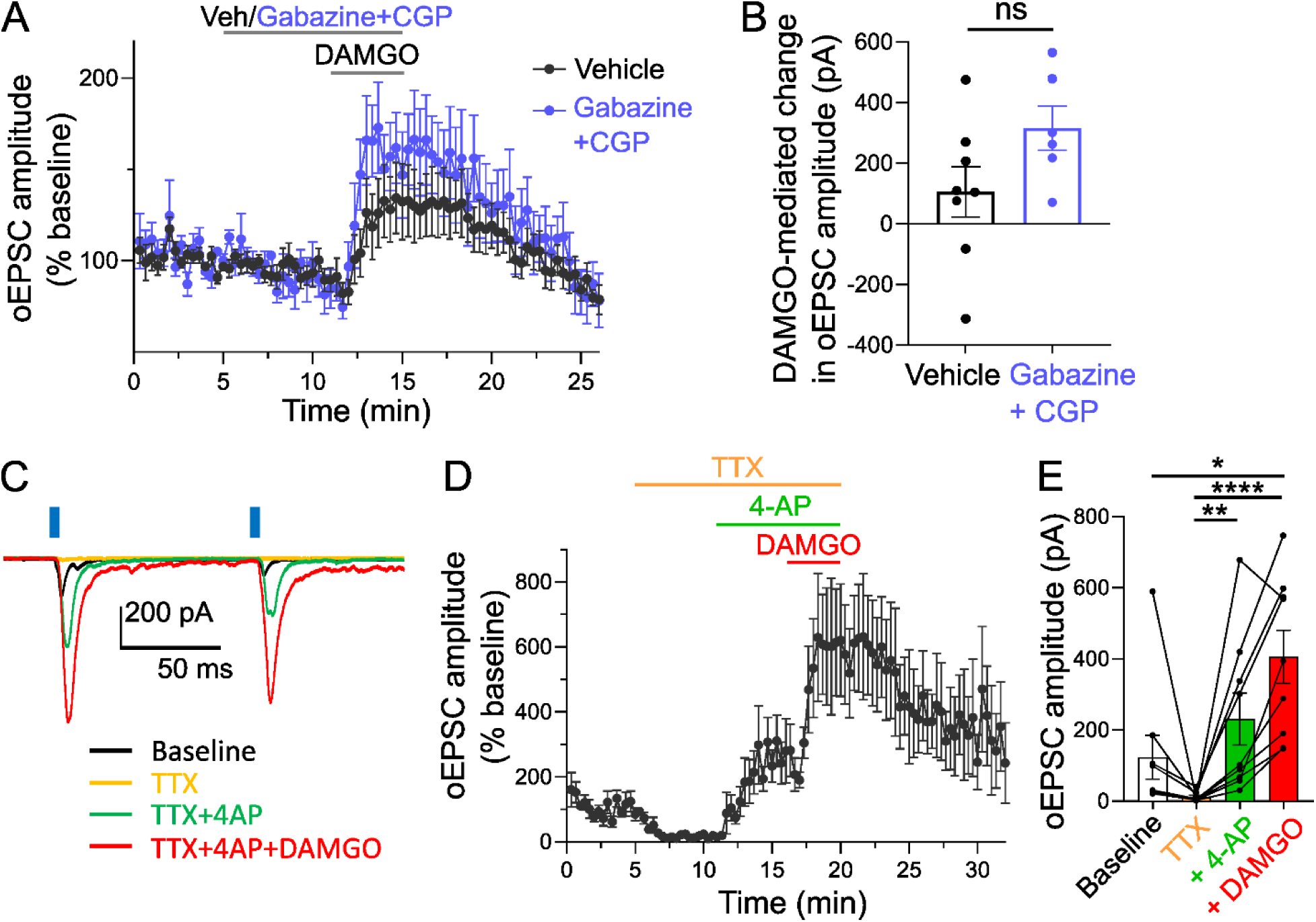
MOR-mediated potentiation of excitatory transmission at the HP synapse persists in the presence of GABA receptor blockers and TTX+4AP. **A**) Averaged time-trace showing DAMGO (1 μM)-mediated increase in oEPSC amplitude at the HP synapse after pre-treatment with gabazine (5 μM) plus CGP (2 μM) (n=6) or vehicle (0.04 % DMSO in ACSF) (n=8). **B**) Effects of DAMGO on oEPSC amplitude persisted in the presence of gabazine plus CGP; unpaired t-test, p= 0.09. **C**) Example trace, **D**) averaged time trace (n=6) and **E**) bar graph (n=9) show DAMGO potentiation of oEPSC amplitude persisted in the presence of TTX+4-AP; Friedman test, *p< 0.05, **p< 0.01, ****p< 0.0001.

**Figure 4.**
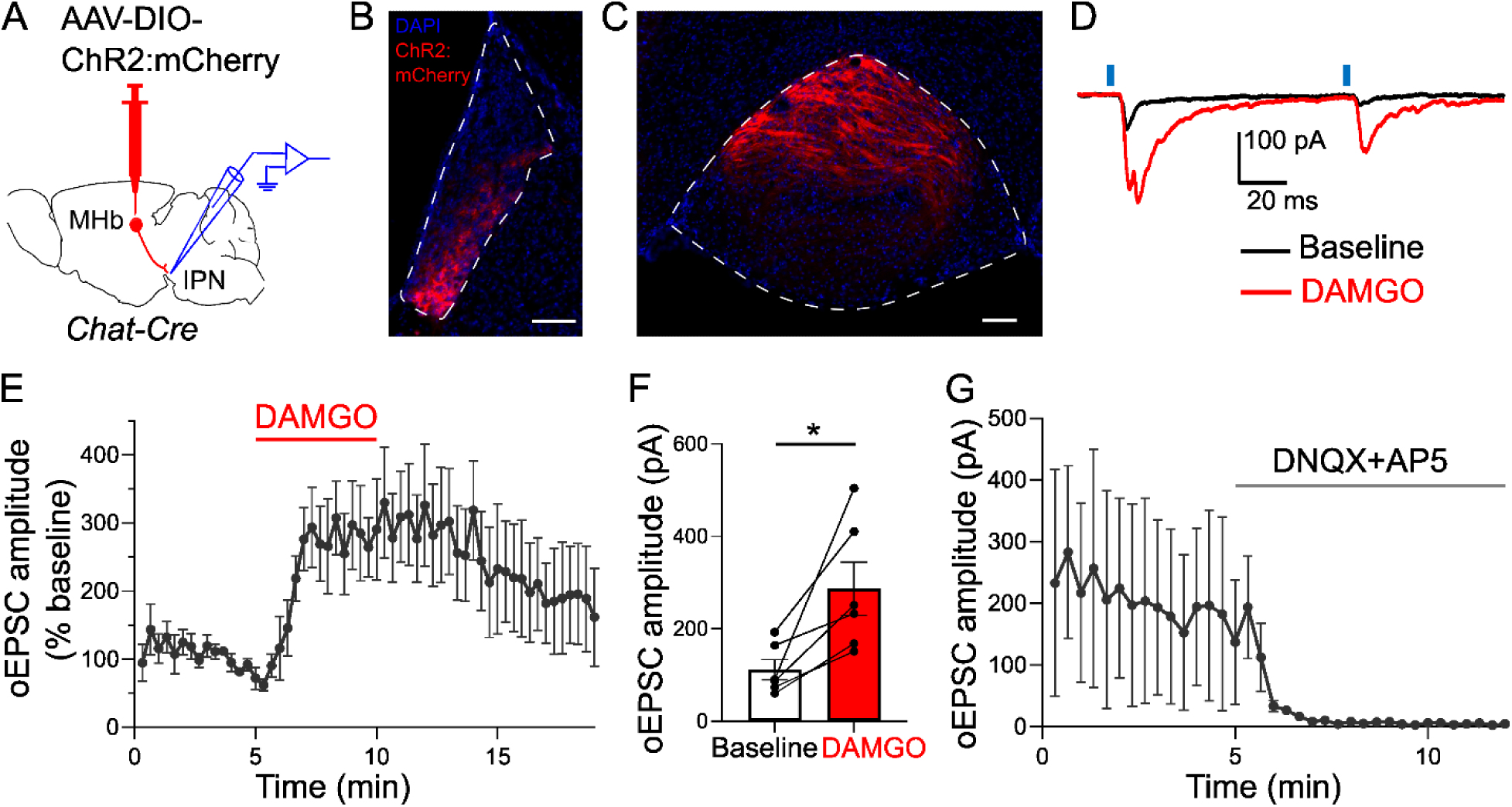
Glutamate co-release at cholinergic HP synapses is potentiated by MOR activation. **A)** Experiment design showing patch-clamp electrophysiology in IPN following AAV injection into MHb of *Chat*-Cre mice. **B**) Images showing ChR2:mCherry expression in ventral MHb and **C**) projections in IPR; scales, 100 μm. **D**) Example traces, **E**) averaged time-trace, and **F**) bar graph show DAMGO (1 μM) potentiation of oEPSC amplitudes at HP cholinergic synapses (n=6); paired t-test, *p<0.05. **G**) Application of glutamate receptor blockers, DNQX (10 μM) plus AP5 (40 μM) abolished oEPSCs in IPR neurons (n=2).

To directly test if the potentiating effects of MOR activation on excitatory neurotransmission are monosynaptic, we first pharmacologically isolated the HP synapse by applying the voltage-gated sodium channel blocker tetrodotoxin (TTX) to prevent the propagation of action potentials, and then used the potassium channel blocker 4-aminopyridine (4-AP) to enhance the effects of optogenetic stimulation on MHb terminals. As before, oEPSCs were evoked in IPR cells under baseline conditions, followed by bath application of TTX which effectively abolished these oEPSCs, then functional recovery of oEPSCs with the application of 4-AP. In these now monosynaptically isolated oEPSCs we applied DAMGO, and found that MOR activation again led to a significant potentiation of oEPSC amplitude (**Figure 3C-E**) (p< 0.0001, Friedman test).

In this experiment we noted that one cell showed an apparent DAMGO-mediated inhibition of neurotransmission (**Figure 3E**). In sum, and across the experiments presented, we found three cells which showed a DAMGO-mediated inhibition rather than potentiation of neurotransmission. We mapped the location of these cells and found that all three were located in caudal IPR (**Supplemental Figure 3D**), suggesting that a subset of caudal IPR cells may not receive input from MOR-expressing MHb axons that show this non-canonical potentiation, but instead from a population of MOR-expressing axons that show canonical inhibitory effects on neurotransmission.

Overall, these results suggest that MOR activation at the HP synapse leads to net potentiation of glutamate transmission in a large majority of IPR cells through a monosynaptic mechanism.

### MOR activation potentiates excitatory transmission from HP cholinergic synapses

The potentiating effects of MOR activation on excitatory transmission are novel. However, activation of GABA_B_ receptors, another G_i_-coupled GPCR were prior shown to potentiate excitatory neurotransmission from cholinergic terminals at HP synapses (Zhang et al., 2016). We thus evaluated the effects of the GABA_B_ receptor agonist, baclofen, on neurotransmission at the HP synapse in MOR-Cre mice. Similar to DAMGO, baclofen appeared to potentiate oEPSC amplitude, at least in a subset (3/4) IPR cells (t_3_= 1.5, p= 0.24) (**Supplemental Figure 4A-E**). These data suggest that MOR and GABA_B_ potentiate excitatory transmission through similar mechanisms from an overlapping population of HP synapses.

Indeed, our RNAscope data (**Supplemental Figure 1A-B**) demonstrates that a subset of cholinergic neurons located in ventrolateral MHb subareas express MOR. To test whether excitatory transmission from IPN-projecting cholinergic neurons is potentiated by MOR activation we expressed ChR2:mCherry in MHb of *Chat*-Cre mice (**Figure 4A**). ChR2:mCherry expression was restricted to ventral regions of MHb (**Figure 4B**) and observed throughout IPR (**Figures 4C**), a pattern consistent with cholinergic projections in the IPN (Ren et al., 2011; Zhang et al., 2016; Souter et al., 2022). Using whole-cell patch clamp we recorded fast oEPSCs in IPR that were significantly potentiated by DAMGO (**Figure 4D-F**) (t_5_= 3.3, p= 0.02). Following DAMGO washout, we applied glutamate blockers in a subset of cells (n=2) which abolished the oEPSCs (**Figure 4G**), indicating they were dependent on glutamate release. Therefore, MHb cholinergic neurons co-release glutamate at the HP synapse, which is enhanced following MOR activation.

### DAMGO potentiates neurotransmission through pre-synaptic MORs at the HP synapse

The above data support the hypothesis that presynaptic MOR activation leads to non-canonical potentiation of excitatory transmission at the HP synapse. To definitively test this hypothesis, we aimed to selectively disrupt MOR expression from presynaptic MHb neurons. To do so, we injected AAV-Cre into MHb of mice carrying a floxed *Oprm1* gene (*Oprm1^fl/fl^*). We also co-injected an AAV for Cre-dependent expression of ChR2:mCherry (**Figure 5A**). This approach will lead to disruption of MOR and simultaneous expression of ChR2:mCherry in Cre-expressing neurons; though ChR2 expression will not be restricted only to MOR-expressing neurons using this approach. We waited >6 weeks to allow for MOR protein turnover before recording oEPSCs in IPR neurons. DAMGO-mediated potentiation of oEPSC amplitude was observed in wild-type (WT) control mice, but not in cells recorded from Oprm1^fl/fl^ mice (**Figure 5B-5D**) (group x treatment interaction: F_1,19_= 11.7, p= 0.003; main effect of treatment: F_1,19_= 2.6, p= 0.12). These results demonstrate that activation of pre-synaptic MOR is required for the non-canonical MOR-induced potentiation of excitatory transmission at the HP synapse.

**Figure 5.**
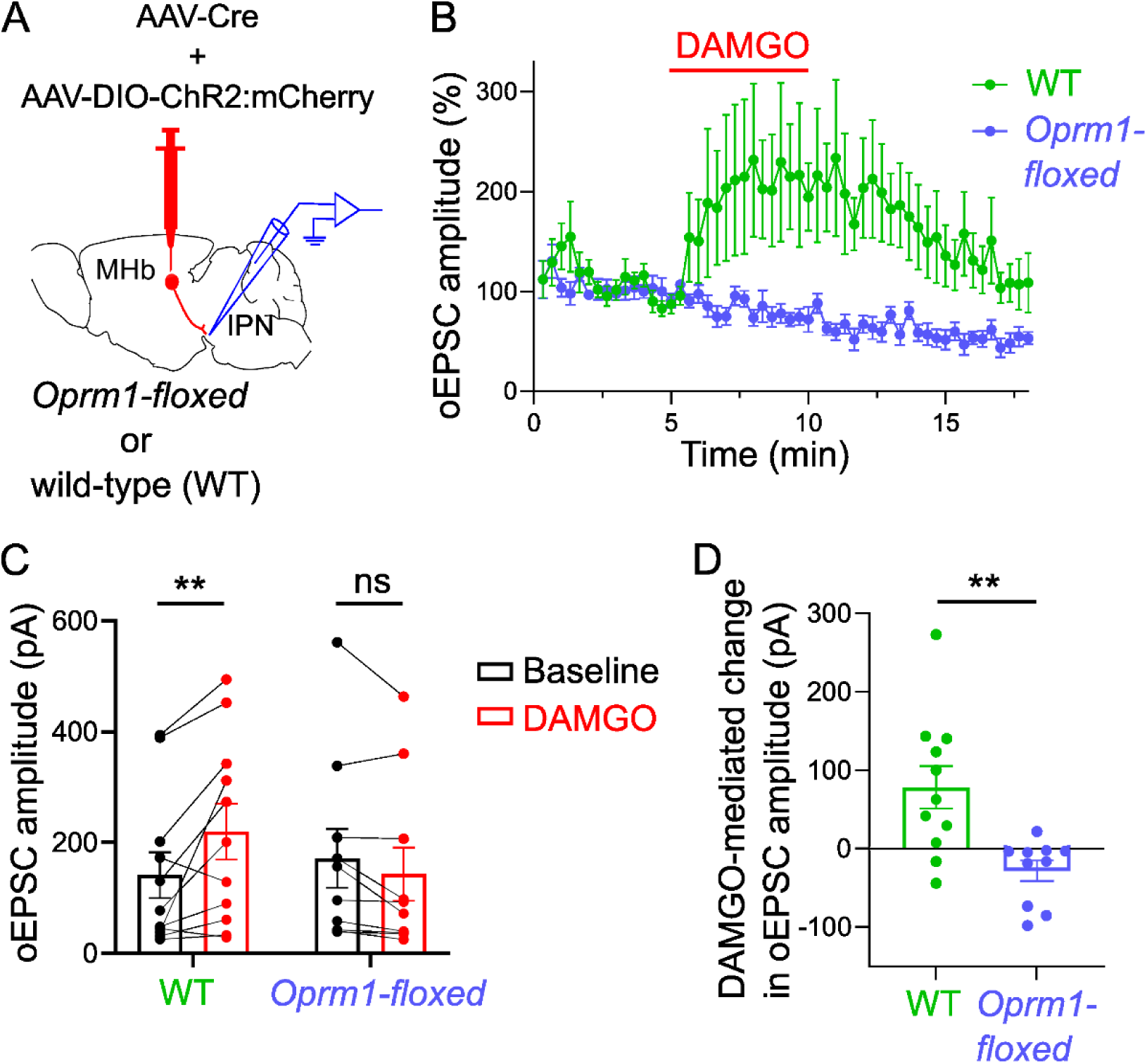
Pre-synaptic MOR is required for DAMGO-mediated potentiation of excitatory transmission at the HP synapse. **A**) Experimental design to delete MOR expression in pre-synaptic MHb neurons. **B**) Averaged time trace and **C**) bar graph show DAMGO (1 μM) potentiation of oEPSC amplitude at the HP synapse in wild-type (WT) mice (n=11), but not *Oprm1^fl/fl^* mice (n=10); mixed-model 2-way ANOVA with Sidak post-hoc multiple comparisons, **p< 0.01. **D**) Significant potentiation of oEPSC amplitude by DAMGO in IPR neurons of WT compared to *Oprm1^fl/fl^* mice; Mann-Whitney test, **p<0.01.

## Discussion

Activity in the HP circuit has been linked to the expression of avoidance and coping behaviors in response to aversive and anxiogenic stimuli, including in the context of addiction (Fowler and Kenny, 2014; Soria-Gómez et al., 2015; Zhang et al., 2016; Klenowski et al., 2023; Molas et al., 2024). Nicotinic acetylcholine receptors (nAChR) are densely expressed in the HP circuit and have been shown to contribute to aversive qualities of nicotine and nicotine-dependence (Salas et al., 2009; Fowler et al., 2011; Antolin-Fontes et al., 2015; McLaughlin et al., 2017; Molas et al., 2017). Nicotine withdrawal increases the activity of IPN neurons and dampening their excitability can alleviate nicotine-withdrawal symptoms (Zhao-Shea et al., 2013, 2015; Klenowski et al., 2022). MORs are also densely expressed in the HP circuit (Mansour et al., 1987; Kitchen et al., 1997). MOR-expressing neurons in MHb contribute to the expression of affective and somatic features of opioid dependence (Boulos et al., 2020). Furthermore, increased IPN activity was observed during opioid withdrawal, while ablation of descending IPN projection neurons reduced the aversive qualities of opioid withdrawal (Liang et al., 2024). Yet there is almost nothing known about how MOR signaling influences the activity of the HP circuit.

In this study we used reporter and conditional knockout mice with optogenetics and patch-clamp electrophysiological recordings to establish both canonical and non-canonical effects of MOR activation on neurons and synapses in the HP circuit. MORs are G_i_- coupled GPCRs and their activation can increase potassium and inhibit calcium conductances, suppressing neuronal activity and neurotransmission (Al-Hasani and Bruchas, 2011; Wang et al., 2019; Birdsong and Williams, 2020; Stoveken et al., 2020; Che and Roth, 2023). Consistent with these canonical mechanisms, we found that within the HP circuit MOR: i) inhibited firing in MOR-expressing MHb neurons, ii) activated an inhibitory hyperpolarizing conductance in IPN neurons, iii) inhibited sEPSC frequency in IPN.

In contrast to these canonical inhibitory effects, MOR activation markedly potentiated evoked glutamatergic transmission at the HP synapse. Through disinhibition MOR activation can produce excitation in several brain regions including hippocampus, periaqueductal gray, and VTA (Charles and Hales, 2004; Henderson, 2015). However, a direct facilitatory effect of MOR activation has not been reported for other synapses. IPN neurons are primarily GABAergic and activation of these neurons can regulate excitatory neurotransmission at the HP synapse through GABA release (Zhang et al., 2016; Koppensteiner et al., 2017). However, the MOR-mediated facilitatory effect on glutamatergic transmission at the HP synapse persisted in the presence of GABA receptor blockade, indicating this effect was not dependent on local GABA signaling. Moreover, the faciliatory effect of MOR persisted when the optogenetic stimulation was monosynaptically isolated, demonstrating that it is not dependent on any other polysynaptic recruitment mechanism. Finally, genetic disruption of MOR in presynaptic MHb neurons completely abolished the faciliatory effect that we detected in postsynaptic IPN neurons. These results indicate that MOR activation on MHb axons has a non-canonical faciliatory effect on glutamate release.

What intracellular mechanisms downstream of MOR activation could potentiate evoked neurotransmitter release? Activation of GABA_B_ receptors, also G_i_-linked GPCRs, has been shown to similarly potentiate excitatory transmission from MHb cholinergic axons in IPN (Zhang et al., 2016; Koppensteiner et al., 2024). Indeed, we provide evidence for a similar effect of GABA_B_ activation from MOR^+^ HP synapses, suggesting overlap in expression and GPCR signaling mechanisms between MOR and GABA_B_ at HP synapses. GABA_B_-mediated potentiation of glutamatergic transmission at the HP synapse was sensitive to pertussis toxin treatment and amplified presynaptic Ca^2+^ entry through Ca_v2.3_ channels (Zhang et al., 2016), but was not dependent on cAMP/PKA or PLC/PKC signaling pathways (Koppensteiner et al., 2024). MOR activation on MHb terminals could therefore be signaling through pertussis toxin sensitive Gi-proteins which non-canonically couple to the same set of Ca^2+^ channels as do the GABA_B_ receptors, thereby enhancing pre-synaptic Ca^2+^ influx and neurotransmitter release at the HP synapse.

Although MOR activation potentiated evoked glutamatergic transmission in a large majority of cells across rostral to caudal IPR, an inhibitory effect was observed in a minority of IPR neurons. Each of these inhibitory responses were detected in caudal IPR neurons. One possibility is that some of the recorded neurons lie at the interface of caudal IPR and the rostral edge of median raphe, and these receive input from a different population of MOR-expressing neurons in either medial or lateral habenula (LHb) (Quina et al., 2015; Ables et al., 2023; Groos and Helmchen, 2024). Indeed, MOR-Cre-expressing LHb neurons also expressed ChR2 in our preparation, and these cells have been shown to shown to express functional MORs with canonical inhibitory effects (Margolis and Fields, 2016).

An alternative explanation relates to the observation that there are also multiple IPN-projecting cell types within MHb (Gardon et al., 2014; Hashikawa et al., 2020; Wallace et al., 2020; Sylwestrak et al., 2022; van de Haar et al., 2022). We show that MOR potentiates glutamate release from cholinergic neurons localizing to ventral MHb, but that MOR is prominently expressed in lateral and dorsal MHb populations that do not express cholinergic markers. Thus, MOR may potentiate excitatory transmission from cholinergic neurons, but inhibit excitatory transmission at some or all of the non-cholinergic HP synapses. In that event, we may be measuring the net effects of evoked glutamate release from a mix of HP synapses that are canonically inhibited and non-canonically potentiated, though the potentiating effects of MOR are clearly more prominent in our preparation.

Yet in the same postsynaptic IPR cells where we observed pronounced MOR-induced potentiation of evoked excitatory transmission, we detected MOR-induced inhibition in the rate of spontaneous excitatory events. This may indicate that MOR activation simultaneously suppresses ‘background’ spontaneous neurotransmission while potentiating evoked neurotransmission from habenular inputs onto IPN cells. Interestingly, activation of GABA_B_ receptors shows the same pattern of differential effects on evoked and spontaneous transmission at the HP synapse (Koppensteiner et al., 2024). These phenomena could arise due to differential GPCR modulation of evoked and spontaneous release arising from distinct pools of vesicles.

Repeated exposure to exogenous opioids can lead to receptor desensitization and tolerance that reduces the number of functional receptors (Williams et al., 2013; Adhikary and Williams, 2022). Opioid-induced changes in MOR signaling within the HP circuit have not been assessed, but MOR in the HP circuit can contribute to the manifestation of behaviors linked to opioid dependence and withdrawal (Boulos et al., 2020; Liang et al., 2024). Future work must therefore investigate how chronic exposure to exogenous opioids changes the HP circuit, and it will be important to assess how MOR signaling changes within distinct cellular and subcellular compartments.

In this study we report the physiological consequences of MOR activation in the HP circuit. Canonical inhibitory effects were observed in somatodendritic compartments of MHb and IPN neurons. However, we also revealed a novel faciliatory effect of MOR on evoked neurotransmission at the HP synapse. These results point to a new mode by which MOR can signal in presynaptic terminals, and provides a new target through which opioids may act to induce neuroadaptations that underlie opioid addiction.

## Author Contributions

Conceptualization, S.M.S., W.S.C., and T.S.H.; Methodology, S.M.S., A.S., Y.C., and T.S.H.; Investigation, S.M.S., A.S., and Y.C.; Formal analysis, S.M.S., A.S., and Y.C.; Writing – original draft, S.M.S. and T.S.H.; Writing – review and editing, S.M.S., A.S., Y.C., W.S.C., and T.S.H.; Funding acquisition, W.S.C., and T.S.H.

## Acknowledgements

Funding for this work was provided by NIH-NIDA awards R21DA054693 and R01DA060923. We are grateful to Dr. Brigitte Kieffer for providing *Oprm1* (MOR)-Cre mice, and for pAAV vectors generated by Dr. Karl Deisseroth and obtained through Addgene.^1^

## Materials and Methods

### Animals

Mice were bred at University of California San Diego (UCSD) and group housed on a 12-h light/dark cycle with ad libitum access to pelleted food and water. Initial breeders wild-type (C57Bl/ 6J, Stock: 000664), Ai6 ZsGreen reporter (Stock: 007906), Ai14 tdTomato reporter (Stock: 007914), Ai27D ChR2:tdTomato reporter (Stock: 012567), Ai32 ChR2:eYFP reporter (Stock: 012569), *Chat^Cre/Cre^* (Stock: 006410) and *Oprm1^fl/fl^* (Stock: 030074) were obtained from Jackson laboratories. MOR^+/Cre^ mice were initially obtained from Dr. Brigitte Kieffer (McGill University) and maintained back-crossed on to C57Bl/6J (Bailly et al., 2020) Male and female mice were used for all experiments in approximately equal proportion, except where noted. All experiments were performed in accordance with protocols approved by UCSD Institutional Animal Care and Use Committee.

### Stereotactic surgeries

For intracranial injections, mice (5–8 weeks) were deeply anesthetized with isoflurane and placed in a stereotaxic apparatus (Kopf Instruments). For optogenetic experiments testing effects of DAMGO in MOR-Cre or *Chat-Cre* mice, bilateral injections (150 nl/ hemisphere) of AAV5-EF1a-DIO-ChR2:mCherry (2-2.5 x 10^13^ gc/ml, Addgene 20297) were made into MHb. For optogenetic experiments testing effects of baclofen in MOR-Cre, bilateral injections (150 nl/ hemisphere) of AAV5-EF1a-DIO-ChR2:eYFP (1.8 x 10^13^ gc/ml, Addgene 20298) were made into MHb. For experiments testing effects of DAMGO in *Oprm1-floxed* or wild-type (WT) mice, AAV5-hSyn-Cre (Addgene 105553) was combined with AAV5-Ef1a-DIO-ChR2:mCherry. Bilateral injections (150 nl/ hemisphere) of this mixture were made into MHb at final concentrations of 1.4-5.3 x 10^12^ gc/ml (AAV-Cre) and 1.9 x 10^13^ gc/ml (AAV-DIO-ChR2:mCherry). Injections into MHb were made at a 20° medio-lateral angle at coordinates (mm relative to bregma): ML= ±1.09, AP= -1.58, DV= −2.56. Injections were made using pulled glass pipettes (∼25 μm aperture diameter) and a Nanoject 3 (Drummond Scientific) at a rate of 10 nl/s with 1-s pulse and 5-s inter-pulse interval. The glass pipettes were held in place for 5 min after infusion before retracting. Animals were treated with a topical antibiotic and administered with the analgesic carprofen (5 mg/kg s.c.; Rimadyl). MOR-Cre or *Chat-Cre* mice were allowed to recover for >3 weeks, whereas *Oprm1-floxed* and WT mice were allowed to recover for >6 weeks before proceeding.

### Patch-clamp electrophysiology

Mice were anesthetized on the day of experiments with an intraperitoneal (IP) injection of sodium pentobarbital (200 mg/kg, Virbac) and transcardially perfused with 15-20 ml of ice-cold oxygenated N-methyl D-glucamine (NMDG)-artificial cerebrospinal fluid (ACSF) containing (in mM): 92 NMDG, 2.5 KCl, 1.25 NaH_2_PO_4_, 30 NaHCO_3_, 20 HEPES, 25 D-glucose, 2 thiourea, 5 Na-ascorbate, 3 Na-pyruvate, 0.5 CaCl_2_ and 10 MgSO_4_. Animals were decapitated and brains rapidly extracted then submerged in ice-cold NMDG-ACSF. 200-µm coronal brain slices containing the IPN or MHb were obtained using a Leica VT 1200S vibratome and transferred to a recovery chamber containing 150 mL of preheated (32-34°C), oxygenated NMDG-ACSF. A 2 M Na+ spike-in solution was then added to the recovery chamber in increasing volumes and 5-minute increments to achieve a controlled rate of Na+ reintroduction (Ting et al., 2018), following which slices were transferred to oxygenated ACSF at room temperature containing (in mM): 115 NaCl, 2.5 KCl, 1.23 NaH_2_PO_4_, 2 MgSO_4_, 10 D-glucose, 2 CaCl_2_, 26 NaHCO_3_, 2 thiourea, 5 Na-ascorbate, and 3 Na-pyruvate. Slices were allowed to equilibrate for at least one hour before recordings, following which they were transferred to a recording chamber maintained at ∼31°C and perfused continuously at a rate of 1.5-2 ml/min with oxygenated ACSF that contained (in mM): 125 NaCl, 2.5 KCl, 1.2 NaH_2_PO_4_, 2 MgSO_4_, 12.5 D-glucose, 2 CaCl_2_, and 26 NaHCO_3_. All solutions were saturated with carbogen (95% O_2_ + 5% CO_2_) to maintain a pH of ∼7.3.

For cell attached recordings, brains were extracted from either MOR-Cre x *Rosa26*-tdTomato reporter mice (4-5 weeks) or wild-type (c57) mice (4-6 weeks). Using reporter mice, MOR^+^ neurons in MHb were visualized through epifluorescence imaging with a 40X water-immersion objective and a Zeiss filter set, and visually guided patch-clamp recordings were made using infrared-differential interference contrast (IR-DIC) illumination (Axiocam MRm, Examiner.A1, Zeiss). Wild-type mice were used to make patch-clamp recordings from un-tagged neurons in lateral MHb. Cell-attached recordings were made using 5-7 MΩ patch pipettes filled with a K^+^ gluconate-based internal solution containing (in mM): 130 K-gluconate, 2 KCl, 2 MgCl_2_, 2 Na_2_-ATP, 0.3 Na-GTP, 10 phosphocreatine, 10 HEPES and 0.2 EGTA. Patch pipettes were prepared from thin walled (1.5 mm/ 1.1 mm) borosilicate glass capillaries (Sutter Instruments) using a micro-pipette puller (Narishige, PC-10). Cell-attached recordings were made in voltage-clamp mode with command voltage set so holding current was approximately zero. Baseline firing was recorded for 5 minutes followed by 5 minutes of DAMGO (5 µM) application then 5 minutes of DAMGO (5 µM) + Naloxone (5 µM) application. The firing frequency was calculated as the average of the last 3 minutes of each condition.

For whole-cell recordings and optogenetic experiments, brain slices were obtained from adult mice (8-15 weeks). Using IR-DIC, patch-clamp recordings were made from IPR cells adjacent to mCherry- or eYFP-positive fibers which were visualized with epifluorescence imaging using a 40X water-immersion objective and a Zeiss filter set. Whole-cell voltage-clamp recordings were made with patch pipettes pulled to an open tip resistance of 6-7 MΩ and filled with either a K-gluconate internal solution (as described above) or a cesium-based internal solution containing (in mM): 130 CsMeSO_3_, 3 MgCl_3_, 4 Na_3_-ATP, 0.25 Na-GTP, 10 phosphocreatine, 10 HEPES, 0.2 EGTA, and 5 QX-314. Internal solutions were adjusted to a pH of ∼7.3 and ∼285mOsm and used consistently within an experiment. To record optogenetic-induced excitatory postsynaptic currents (oEPSCs), cells were voltage-clamped at -65 mV and two pulses of blue light (2 or 5 ms) were flashed at 10 Hz through the light path of the microscope using a light-emitting diode (UHP-LED460, Prizmatix) under computer control to activate ChR2. The paired light pulses were delivered every 20 seconds. In all experiments, oEPSCs were recorded for a baseline period of 5 minutes, DAMGO (1 µM) was bath applied for either 4 or 5 minutes, followed by washout. Light-pulse width and DAMGO application duration was kept consistent within an experiment. Changes in amplitude of the first oEPSC were averaged and reported in graphs. Baseline oEPSC amplitude was averaged from the last 3 minutes of baseline recording; the oEPSC amplitude in response to DAMGO was averaged from the last 2 or 3 minutes of 4- or 5-minute DAMGO application. Glutamate receptor blockers DNQX (10 µM) and AP5 (40 µM) were bath applied ∼10 minutes after DAMGO washout to test effects of glutamate receptor blockade on oEPSCs. For experiments testing effects of DAMGO with Gabazine and CGP, after baseline recording of oEPSCs, Gabazine (5 µM) and CGP-55845 (2 µM) was bath applied for 6 minutes followed by application of DAMGO + Gabazine + CGP for 4 minutes. For control experiments, vehicle (0.04 % DMSO in ACSF) was applied instead of Gabazine+CGP. For experiments testing effects of DAMGO with picrotoxin (PTX) and CGP, after baseline recording of oEPSCs, PTX (50 µM) and CGP (2 µM) was bath applied for 10 minutes followed by application of DAMGO + PTX + CGP for 5 minutes. For control experiments, vehicle (0.09 % DMSO in ACSF) was applied instead of PTX+CGP. For experiments testing effects of DAMGO with tetrodotoxin (TTX) and 4-aminopyridine (4-AP), after baseline recording of oEPSCs, TTX (1 µM) was bath applied for 6 minutes followed by application of 4-AP (100 µM) + TTX for 5 minutes, followed by application of DAMGO + 4-AP + TTX for 4 minutes. The oEPSC amplitude in response to TTX was averaged from the last 3 minutes of its application before 4-AP, and the oEPSC amplitude in response to 4-AP was averaged from the last 2 minutes of its application before DAMGO. TTX and 4-AP application duration was different in 3 out of 9 cells and therefore not included in the averaged time-trace graph. For testing the effects of baclofen on oEPSC amplitude, baclofen (5 µM) was bath applied for 5 minutes after baseline recording and the oEPSC amplitude was averaged from the last 3 minutes of its application. The holding current was extracted from the last 10 seconds of the 20 second sweep from cells which were patched with the K-gluconate internal solution. Cells with a rate of change of holding current (dI/dt)> 10 pA/ min after DAMGO application were categorized as responding cells. Changes in the frequency of spontaneous excitatory post-synaptic currents (sEPSCs) were also monitored in a subset of these cells in response to DAMGO application. Baseline and DAMGO holding current was averaged from the last 3 minutes of baseline recording and DAMGO application, respectively. Baseline sEPSC frequency was averaged from the last 3 minutes of baseline recording, whereas during DAMGO, was averaged from a 2-minute window, 2 minutes after start of DAMGO application. Experiments and data analysis for testing effects of DAMGO on oEPSCs in WT or *Oprm1^fl/fl^* mice were performed by experimenter blind to genotype. Access resistance (Ra) was calculated from a 10-mV hyperpolarizing pulse applied at the beginning of each sweep and monitored for whole-cell recordings. Cells with a Ra change> 30% were excluded from analyses.

Recordings were made using a Multiclamp 700B Amplifier (Axon Instruments), lowpass filtered at 2 kHz, digitized at 20 kHz (Axon Digidata 1550, Axon Instruments), collected and analyzed using Clampex and Clampfit v10 software (Molecular Devices). All drugs were added to the ACSF following dilution of a stock solution stored at -20°C. All drugs were purchased from Tocris, except PTX, which was purchased from Sigma-Aldrich.

### RNAscope

Adult female wild-type mice (10 and 19 weeks) were anesthetized with an IP injection of sodium pentobarbital (200 mg/kg) and transcardially perfused with 20 ml UltraPure water-based 1x PBS (water from Invitrogen; 10x PBS from Fisher Bioreagents) followed by 20 ml 4% PFA at a rate of 6 ml/min. Brains were extracted, fixed in 4% PFA overnight in 4° C and transferred to a 30% sucrose solution (in PBS) for 24h at 4 °C. 20- µm-thick coronal sections containing MHb or IPN were made using a Leica CM3050S cryostat and slices were collected in PBS and mounted on Superfrost Plus microscope slides (Fisher Scientific). RNAscope *in-situ* fluorescent hybridization was done as previously published (Flores and Hnasko, 2024) using probes (Advanced Cell Diagnostics) against *Oprm1*, *Chat,* and *Slc17a6* (VGLUT2). Four mice were sacrificed to obtain MHb or IPN sections, which were imaged with an Axio Observer.Z1 fluorescent microscope using ApoTome.2 system and Plan-APOCHROMAT 20x/0.8 objective (Zeiss).

### Fluorescent imaging

Native fluorescence was imaged from sections expressing ZsGreen or mCherry. Mice were anesthetized with an IP injection of sodium pentobarbital (200 mg/kg; i.p.) and transcardially perfused with 20 ml of phosphate-buffered saline (PBS) followed by 20 ml 4% paraformaldehyde (PFA) at a rate of 6 ml/min. Brains were extracted, post-fixed in 4% PFA at 4° C overnight, and transferred to 30% sucrose in PBS for 48 h at 4° C. Brains were flash frozen in isopentane and stored at -80° C. 30-μm coronal sections containing the MHb and IPN were obtained using a cryostat (CM3050S, Leica) and collected in PBS containing 0.01% sodium azide. Sections were then mounted on slides, and coverslipped with Fluoromount-G mounting medium (Southern Biotech) containing DAPI (0.5 μg/ml; Roche). Images were acquired using widefield epifluorescence (Zeiss AxioObserver) with a 10X and 63X objective using ApoTome.2 system. MHb subnuclei borders were drawn based on (Juárez-Leal et al., 2022). IPN subnuclei borders were drawn based on “The Mouse Brain in Stereotaxic Coordinates” (Paxinos, G. and Franklin, K.B.J. (2001) The Mouse Brain in Stereotaxic Coordinates. 2nd Edition, Academic Press).

### Statistical analysis

Statistical analysis was performed using GraphPad Prism v10. All data are represented as mean ± standard error of the mean (SEM) and/or as individual points. Data was tested for normality using the Shapiro-Wilk test and parametric or non-parametric tests were used as appropriate. Data was analyzed using paired t-test or Wilcoxon test, unpaired t-test or Mann-Whitney test, Friedman test and 2-way repeated-measures ANOVA followed by Sidak post-hoc multiple comparisons. Details of statistical tests used and sample sizes, can be found in results and in figure legends.

## Figures

**Supplemental Figure 1 (related to Figure 1).**
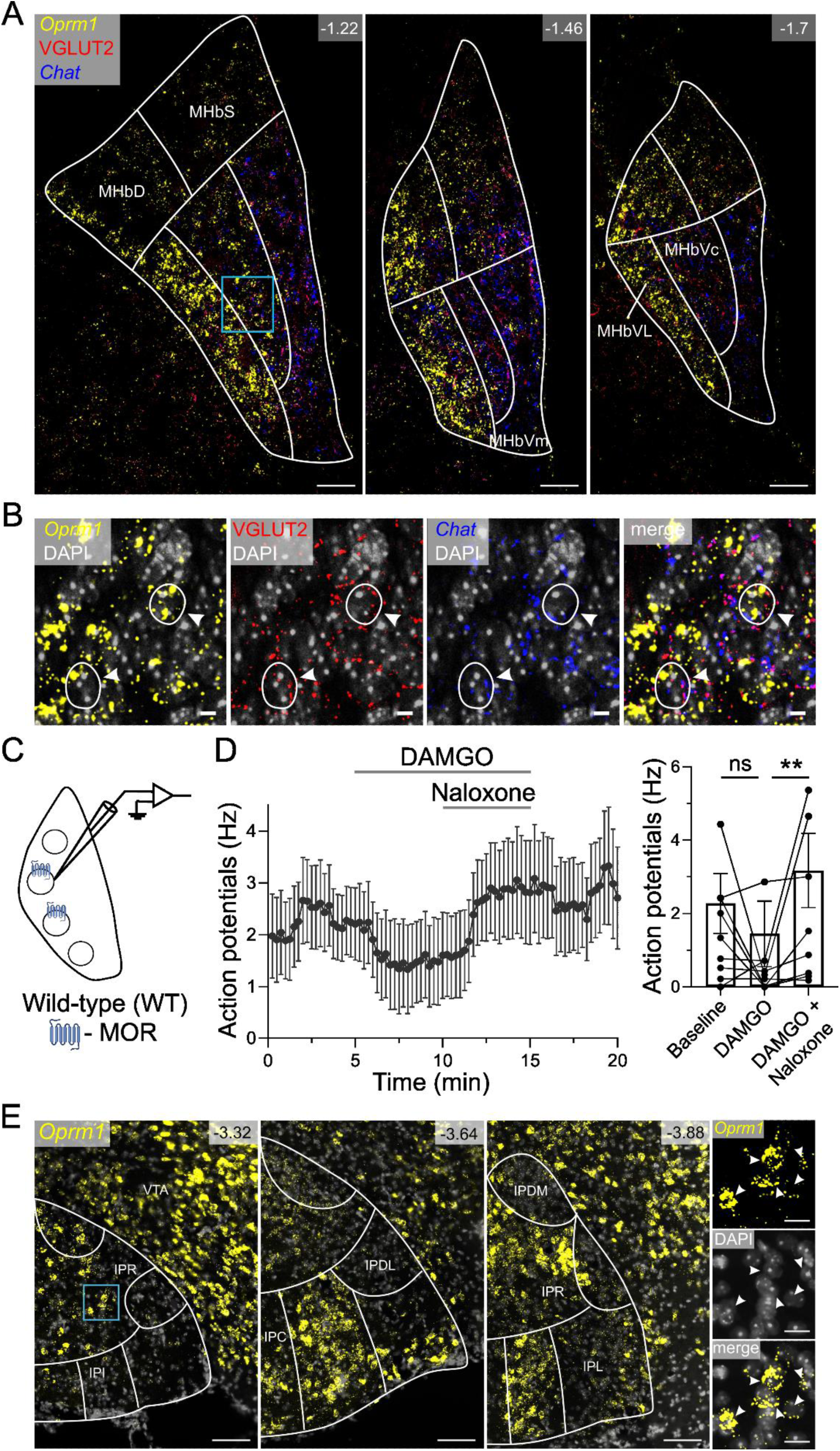
MOR expression in MHb and IPN. **A**) MHb labeled using probes targeting *Oprm1* (MOR, yellow), *Chat* (blue), and *Slc17a6* (VGLUT2, red); coordinates noted relative to bregma (in mm); scale, 50 µm. **B**) Magnified image from inset in panel A (marked by blue square). Arrowheads and outlines indicate MHb neurons co-expressing *Oprm1*, *Chat* and VGLUT2; scale, 20 µm. **C**) Schematic of cell-attached recordings from unlabeled/unidentified lateral MHb cells, some of which express MORs. **D**) Averaged time-trace (left) and bar-graph (right) of cell-attached recordings showing effects of DAMGO (5 μM) and naloxone (5 μM) on action-potential firing in MHb neurons (n=10); Friedman test, *p<0.05. DAMGO application inhibited firing in a subset of neurons (n=4/10), which was reversed by naloxone, consistent with MOR expression in a subset of neurons. **E**) IPN labeled using probes targeting *Oprm1* (MOR, yellow); coordinates noted relative to bregma (in mm); scale, 100 µm.. Right panel set shows magnified area within blue square; arrowheads indicate IPN neurons expressing *Oprm1*; scale, 20 µm.

**Supplemental Figure 2 (related to Figure 2).**
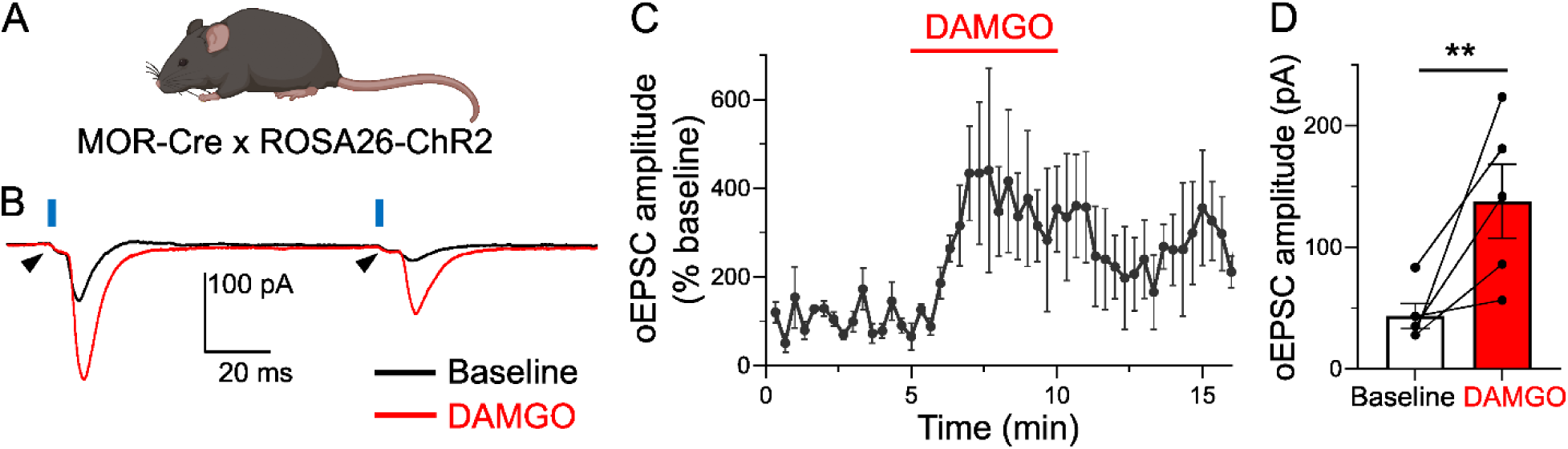
MOR activation potentiates excitatory neurotransmission in the IPR. **A**) MOR-Cre crossed to Ai27 or Ai32 mice resulting in Cre-dependent expression of ChR2:mCherry or ChR2:YFP in cells that expressed MOR. Example traces from IPR neuron showing oEPSCs before and after DAMGO (1 μM) application. Black arrowhead indicates photocurrent, suggestive of a MOR^+^ postsynaptic IPR neuron expressing ChR2. **C**) Average time trace and **D**) bar graph show DAMGO-mediated potentiation of oEPSC amplitude in IPR neurons (n=5); paired t-test, *p< 0.05.

**Supplemental Figure 3 (related to Figure 3).**
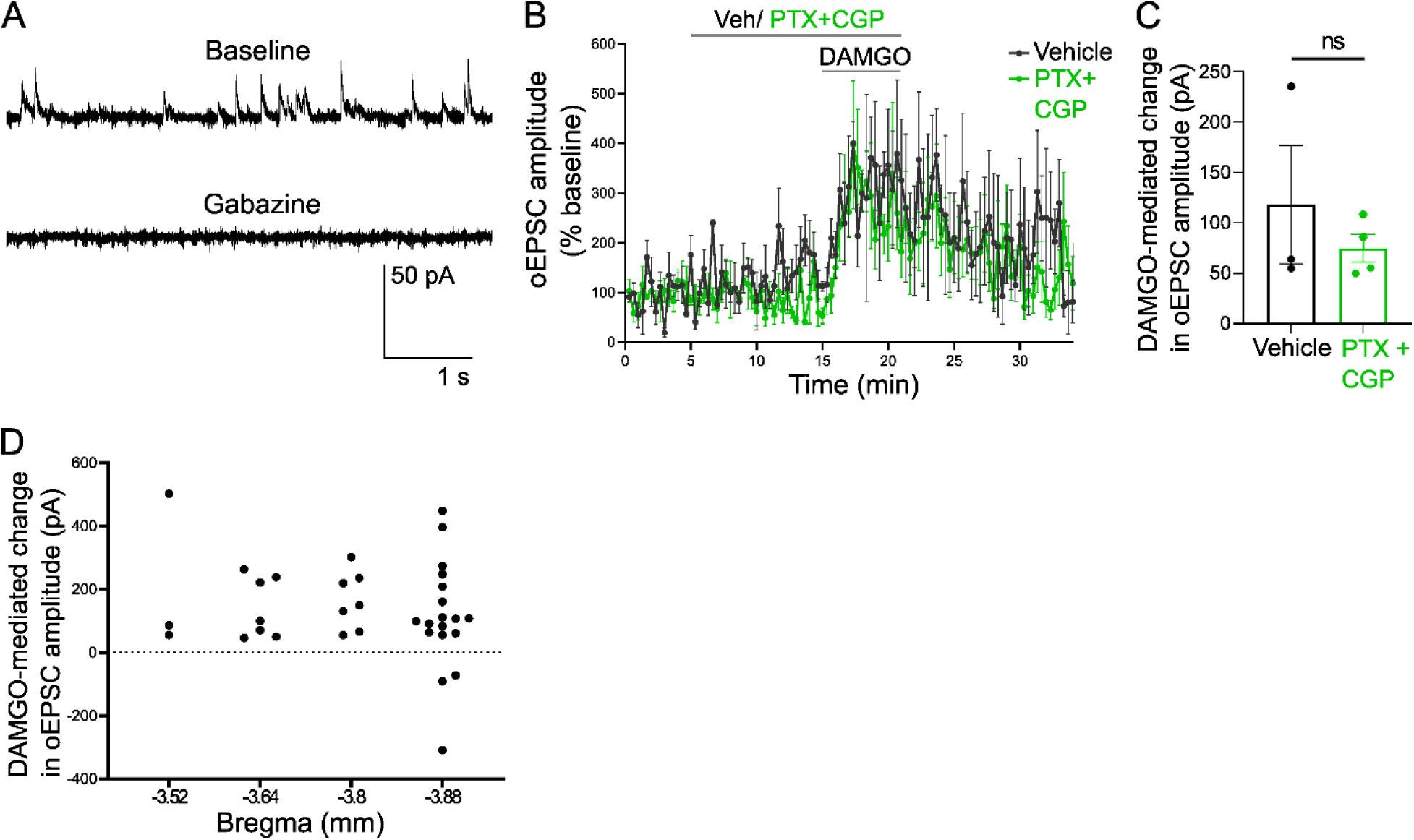
Non-canonical potentiation of excitatory transmission by MOR persists in the presence of GABA blockers, and are observed in IPR cells across the rostral caudal axis, though some caudal IPR cells show canonical inhibition. **A**) Example trace showing sIPSCs in an IPR neuron are blocked by gabazine (5 μM). **B**) Averaged time-trace and (**C**) bar graph showing DAMGO (1 μM) potentiation of oEPSC amplitude (ChR2 expressed in MHb of MOR-Cre mice) persisted in the presence of PTX (50 μM) plus CGP (2 μM) (n=4) or vehicle (n=3); unpaired t-test, p= 0.44. **D**) MOR potentiation of excitatory transmission was observed across the rostral-caudal extent of IPR. However, a subset of IPR neurons in caudal IPR appeared to show a canonical inhibitory effect of MOR activation. These data are a re-representation of DAMGO responses from across the experiments shown in Figures 2, 3 and S3.

**Supplemental Figure 4.**
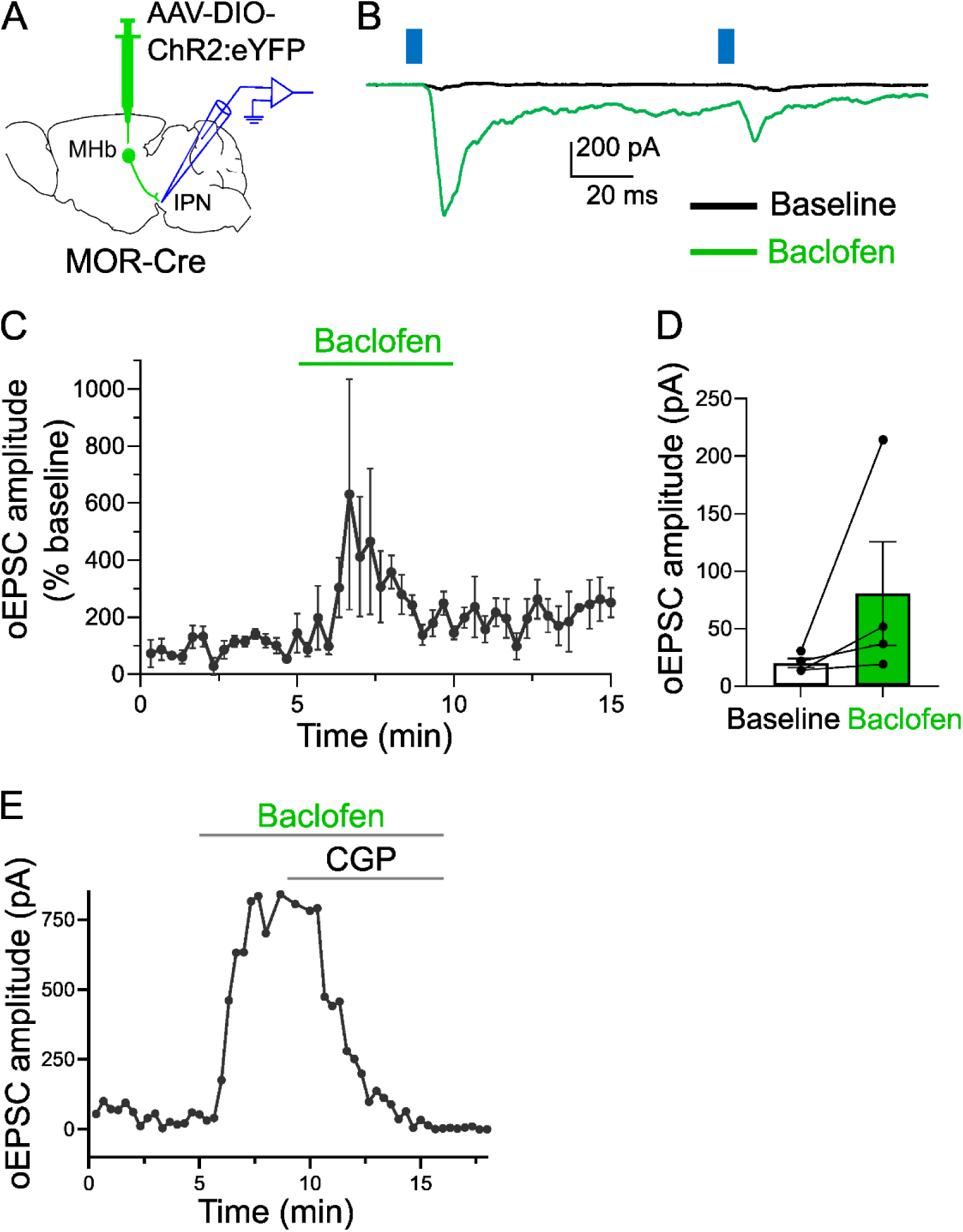
GABAB receptor activation can potentiate excitatory transmission at the MOR^+^ HP synapse. **A**) Experiment design showing patch-clamp electrophysiology in IPN following AAV injection into MHb of MOR-Cre mice. **B**) Example trace showing baclofen (5 μM) potentiation of oEPSC amplitude in an IPR neuron. **C**) Averaged time trace and **D**) bar graph show baclofen-mediated change in oEPSC amplitude (n=4); paired t-test, p= 0.24. **E**) Example time trace from IPR neuron showing baclofen (5 μM) potentiation of oEPSCs, reversed by the GABA_B_ receptor antagonist CGP-55845 (2 μM). Two points were truncated because oEPSCs led to unclamped action-potentials.

## Footnotes

1 This bioRxiv submission was coordinated with a similar study from the McBain laboratory (NIH) entitled ‘Complex opioid driven modulation of glutamatergic and cholinergic neurotransmission in a GABAergic brain nucleus associated with emotion, reward and addiction’ by Chittajallu *et al*.

## References

Ables JL, Park K, Ibañez-Tallon I (2023) Understanding the habenula: A major node in circuits regulating emotion and motivation. Pharmacol Res 190:106734.

Adhikary S, Williams JT (2022) Cellular Tolerance Induced by Chronic Opioids in the Central Nervous System. Front Syst Neurosci 16:937126.

Al-Hasani R, Bruchas MR (2011) Molecular mechanisms of opioid receptor-dependent signaling and behavior. Anesthesiology 115:1363–1381.

Allain F, Carter M, Dumas S, Darcq E, Kieffer BL (2022) The mu opioid receptor and the orphan receptor GPR151 contribute to social reward in the habenula. Sci Rep 12:20234.

Antolin-Fontes B, Ables JL, Görlich A, Ibañez-Tallon I (2015) The habenulo-interpeduncular pathway in nicotine aversion and withdrawal. Neuropharmacology 96:213–222.

Bailly J, Allain F, Schwartz E, Tirel C, Dupuy C, Petit F, Diana MA, Darcq E, Kieffer BL (2023) Habenular Neurons Expressing Mu Opioid Receptors Promote Negative Affect in a Projection-Specific Manner. Biol Psychiatry 93:1108–1117.

Bailly J, Del Rossi N, Runtz L, Li J-J, Park D, Scherrer G, Tanti A, Birling M-C, Darcq E, Kieffer BL (2020) Targeting Morphine-Responsive Neurons: Generation of a Knock-In Mouse Line Expressing Cre Recombinase from the Mu-Opioid Receptor Gene Locus. eNeuro 7:ENEURO.0433-19.2020.

Birdsong WT, Williams JT (2020) Recent Progress in Opioid Research from an Electrophysiological Perspective. Mol Pharmacol 98:401–409.

Boulos LJ, Ben Hamida S, Bailly J, Maitra M, Ehrlich AT, Gavériaux-Ruff C, Darcq E, Kieffer BL (2020) Mu opioid receptors in the medial habenula contribute to naloxone aversion. Neuropsychopharmacol Off Publ Am Coll Neuropsychopharmacol 45:247–255.

Charles AC, Hales TG (2004) From inhibition to excitation: functional effects of interaction between opioid receptors. Life Sci 76:479–485.

Che T, Roth BL (2023) Molecular basis of opioid receptor signaling. Cell 186:5203–5219.

Flores AJ, Hnasko TS (2024) RNAscope. Available at: https://www.protocols.io/view/v2-rnascope-c7vbzn2n [Accessed November 17, 2024].

Fowler CD, Kenny PJ (2014) Nicotine aversion: Neurobiological mechanisms and relevance to tobacco dependence vulnerability. Neuropharmacology 76 Pt B:533–544.

Fowler CD, Lu Q, Johnson PM, Marks MJ, Kenny PJ (2011) Habenular α5 nicotinic receptor subunit signalling controls nicotine intake. Nature 471:597–601.

Gackenheimer SL, Suter TM, Pintar JE, Quimby SJ, Wheeler WJ, Mitch CH, Gehlert DR, Statnick MA (2005) Localization of opioid receptor antagonist [3H]-LY255582 binding sites in mouse brain: comparison with the distribution of mu, delta and kappa binding sites. Neuropeptides 39:559–567.

Gardon O, Faget L, Chu Sin Chung P, Matifas A, Massotte D, Kieffer BL (2014) Expression of mu opioid receptor in dorsal diencephalic conduction system: new insights for the medial habenula. Neuroscience 277:595–609.

Groos D, Helmchen F (2024) The lateral habenula: A hub for value-guided behavior. Cell Rep 43:113968.

Hashikawa Y, Hashikawa K, Rossi MA, Basiri ML, Liu Y, Johnston NL, Ahmad OR, Stuber GD (2020) Transcriptional and Spatial Resolution of Cell Types in the Mammalian Habenula. Neuron 106:743–758.e5.

Henderson G (2015) The μ-opioid receptor: an electrophysiologist’s perspective from the sharp end. Br J Pharmacol 172:260–267.

Juárez-Leal I, Carretero-Rodríguez E, Almagro-García F, Martínez S, Echevarría D, Puelles E (2022) Stria medullaris innervation follows the transcriptomic division of the habenula. Sci Rep 12:10118.

Kitchen I, Slowe SJ, Matthes HW, Kieffer B (1997) Quantitative autoradiographic mapping of mu-, delta- and kappa-opioid receptors in knockout mice lacking the mu-opioid receptor gene. Brain Res 778:73–88.

Klenowski PM, Zhao-Shea R, Freels TG, Molas S, Tapper AR (2022) Dynamic activity of interpeduncular nucleus GABAergic neurons controls expression of nicotine withdrawal in male mice. Neuropsychopharmacol Off Publ Am Coll Neuropsychopharmacol 47:641– 651.

Klenowski PM, Zhao-Shea R, Freels TG, Molas S, Zinter M, M’Angale P, Xiao C, Martinez-Núñez L, Thomson T, Tapper AR (2023) A neuronal coping mechanism linking stress-induced anxiety to motivation for reward. Sci Adv 9:eadh9620.

Koppensteiner P, Bhandari P, Önal C, Borges-Merjane C, Le Monnier E, Roy U, Nakamura Y, Sadakata T, Sanbo M, Hirabayashi M, Rhee J, Brose N, Jonas P, Shigemoto R (2024) GABAB receptors induce phasic release from medial habenula terminals through activity-dependent recruitment of release-ready vesicles. Proc Natl Acad Sci U S A 121:e2301449121.

Koppensteiner P, Melani R, Ninan I (2017) A Cooperative Mechanism Involving Ca2+-Permeable AMPA Receptors and Retrograde Activation of GABAB Receptors in Interpeduncular Nucleus Plasticity. Cell Rep 20:1111–1122.

Kosten TR, Baxter LE (2019) Review article: Effective management of opioid withdrawal symptoms: A gateway to opioid dependence treatment. Am J Addict 28:55–62.

Liang J, Zhou Y, Feng Q, Zhou Y, Jiang T, Ren M, Jia X, Gong H, Di R, Jiao P, Luo M (2024) A brainstem circuit amplifies aversion. Neuron 112:3634–3650.e5.

Lima LB, Bueno D, Leite F, Souza S, Gonçalves L, Furigo IC, Donato J, Metzger M (2017) Afferent and efferent connections of the interpeduncular nucleus with special reference to circuits involving the habenula and raphe nuclei. J Comp Neurol 525:2411–2442.

Mansour A, Khachaturian H, Lewis ME, Akil H, Watson SJ (1987) Autoradiographic differentiation of mu, delta, and kappa opioid receptors in the rat forebrain and midbrain. J Neurosci Off J Soc Neurosci 7:2445–2464.

Margolis EB, Fields HL (2016) Mu Opioid Receptor Actions in the Lateral Habenula. PloS One 11:e0159097.

McLaughlin I, Dani JA, De Biasi M (2017) The medial habenula and interpeduncular nucleus circuitry is critical in addiction, anxiety, and mood regulation. J Neurochem 142 Suppl 2:130–143.

Molas S, DeGroot SR, Zhao-Shea R, Tapper AR (2017) Anxiety and Nicotine Dependence: Emerging Role of the Habenulo-Interpeduncular Axis. Trends Pharmacol Sci 38:169–180.

Molas S, Freels TG, Zhao-Shea R, Lee T, Gimenez-Gomez P, Barbini M, Martin GE, Tapper AR (2024) Dopamine control of social novelty preference is constrained by an interpeduncular-tegmentum circuit. Nat Commun 15:2891.

Pergolizzi JV, Raffa RB, Rosenblatt MH (2020) Opioid withdrawal symptoms, a consequence of chronic opioid use and opioid use disorder: Current understanding and approaches to management. J Clin Pharm Ther 45:892–903.

Quina LA, Harris J, Zeng H, Turner EE (2017) Specific connections of the interpeduncular subnuclei reveal distinct components of the habenulopeduncular pathway. J Comp Neurol 525:2632–2656.

Quina LA, Tempest L, Ng L, Harris JA, Ferguson S, Jhou TC, Turner EE (2015) Efferent pathways of the mouse lateral habenula. J Comp Neurol 523:32–60.

Ren J, Qin C, Hu F, Tan J, Qiu L, Zhao S, Feng G, Luo M (2011) Habenula “cholinergic” neurons co-release glutamate and acetylcholine and activate postsynaptic neurons via distinct transmission modes. Neuron 69:445–452.

Salas R, Sturm R, Boulter J, De Biasi M (2009) Nicotinic receptors in the habenulo-interpeduncular system are necessary for nicotine withdrawal in mice. J Neurosci Off J Soc Neurosci 29:3014–3018.

Soria-Gómez E, Busquets-Garcia A, Hu F, Mehidi A, Cannich A, Roux L, Louit I, Alonso L, Wiesner T, Georges F, Verrier D, Vincent P, Ferreira G, Luo M, Marsicano G (2015) Habenular CB1 Receptors Control the Expression of Aversive Memories. Neuron 88:306– 313.

Souter EA, Chen Y-C, Zell V, Lallai V, Steinkellner T, Conrad WS, Wisden W, Harris KD, Fowler CD, Hnasko TS (2022) Disruption of VGLUT1 in Cholinergic Medial Habenula Projections Increases Nicotine Self-Administration. eNeuro 9:ENEURO.0481-21.2021.

Stoveken HM, Zucca S, Masuho I, Grill B, Martemyanov KA (2020) The orphan receptor GPR139 signals via Gq/11 to oppose opioid effects. J Biol Chem 295:10822–10830.

Sylwestrak EL, Jo Y, Vesuna S, Wang X, Holcomb B, Tien RH, Kim DK, Fenno L, Ramakrishnan C, Allen WE, Chen R, Shenoy KV, Sussillo D, Deisseroth K (2022) Cell-type-specific population dynamics of diverse reward computations. Cell 185:3568–3587.e27.

Ting JT, Lee BR, Chong P, Soler-Llavina G, Cobbs C, Koch C, Zeng H, Lein E (2018) Preparation of Acute Brain Slices Using an Optimized N-Methyl-D-glucamine Protective Recovery Method. J Vis Exp JoVE:53825.

van de Haar LL, Riga D, Boer JE, Garritsen O, Adolfs Y, Sieburgh TE, van Dijk RE, Watanabe K, van Kronenburg NCH, Broekhoven MH, Posthuma D, Meye FJ, Basak O, Pasterkamp RJ (2022) Molecular signatures and cellular diversity during mouse habenula development. Cell Rep 40:111029.

Viswanath H, Carter AQ, Baldwin PR, Molfese DL, Salas R (2013) The medial habenula: still neglected. Front Hum Neurosci 7:931.

Volkow ND, Jones EB, Einstein EB, Wargo EM (2019) Prevention and Treatment of Opioid Misuse and Addiction: A Review. JAMA Psychiatry 76:208–216.

Wallace ML, Huang KW, Hochbaum D, Hyun M, Radeljic G, Sabatini BL (2020) Anatomical and single-cell transcriptional profiling of the murine habenular complex. eLife 9:e51271.

Wang D, Stoveken HM, Zucca S, Dao M, Orlandi C, Song C, Masuho I, Johnston C, Opperman KJ, Giles AC, Gill MS, Lundquist EA, Grill B, Martemyanov KA (2019) Genetic behavioral screen identifies an orphan anti-opioid system. Science 365:1267–1273.

Welsch L, Bailly J, Darcq E, Kieffer BL (2020) The Negative Affect of Protracted Opioid Abstinence: Progress and Perspectives From Rodent Models. Biol Psychiatry 87:54–63.

Williams JT, Christie MJ, Manzoni O (2001) Cellular and synaptic adaptations mediating opioid dependence. Physiol Rev 81:299–343.

Williams JT, Ingram SL, Henderson G, Chavkin C, von Zastrow M, Schulz S, Koch T, Evans CJ, Christie MJ (2013) Regulation of μ-opioid receptors: desensitization, phosphorylation, internalization, and tolerance. Pharmacol Rev 65:223–254.

Xu C, Sun Y, Cai X, You T, Zhao H, Li Y, Zhao H (2018) Medial Habenula-Interpeduncular Nucleus Circuit Contributes to Anhedonia-Like Behavior in a Rat Model of Depression. Front Behav Neurosci 12:238.

Zhang J, Tan L, Ren Y, Liang J, Lin R, Feng Q, Zhou J, Hu F, Ren J, Wei C, Yu T, Zhuang Y, Bettler B, Wang F, Luo M (2016) Presynaptic Excitation via GABAB Receptors in Habenula Cholinergic Neurons Regulates Fear Memory Expression. Cell 166:716–728.

Zhao-Shea R, DeGroot SR, Liu L, Vallaster M, Pang X, Su Q, Gao G, Rando OJ, Martin GE, George O, Gardner PD, Tapper AR (2015) Increased CRF signalling in a ventral tegmental area-interpeduncular nucleus-medial habenula circuit induces anxiety during nicotine withdrawal. Nat Commun 6:6770.

Zhao-Shea R, Liu L, Pang X, Gardner PD, Tapper AR (2013) Activation of GABAergic neurons in the interpeduncular nucleus triggers physical nicotine withdrawal symptoms. Curr Biol CB 23:2327–2335.

